# Polymer dynamics reveal stage-wise unfolding and homologous pairing of chromosomes in S. pombe

**DOI:** 10.1101/2025.04.30.651601

**Authors:** Sandeep Kumar, Ranjith Padinhateeri, Snigdha Thakur

## Abstract

During meiosis, the exchange of genetic material (“recombination”) is essential to bring up the genetic variation in offspring. To carry out the recombination, homologous pairs of DNA have to come in proximity and align. In fission yeast, this is accomplished by the extensive nuclear oscillation that applies pulling force on the chromosomes. In this work, we investigate the competition between the pulling force that tries to facilitate the alignment and the intra–chromatin interactions that assist the chromatin folding. The crucial role played by the intra–chromatin interaction for a range of pulling forces and loci location is highlighted here. Upon comparing our results with the experimental findings, we not only reaffirm the enhancement of contact probability with loci–dependent pulling but also get a meaningful interpretation of the chromosome unfolding and pairing in different stages of movement.

## I. INTRODUCTION

Chromatin, a long polymer that resides in the nucleus, is made of DNA and proteins and encodes genetic information. When cells divide, genetic information is copied by replicating DNA and reassembling chromatin. In every organism, there are two types of cell division known as mitosis and meiosis. While somatic (non-reproductive) cells undergo mitosis and are responsible for the growth of the organism, gametes (reproductive cells) undergo meiosis and are responsible for life and genetic inheritance [1]. For successful meiosis, in addition to copying the genome (chromosome), an extra process called “crossing over” is necessary. In this process, the pair of chromosomes containing the same genes in the same order (for example, maternal and paternal chromosomes), need to come in close physical proximity (“homologous pairing”) and exchange genetic material (“recombination”) [1–3]. Two homologous chromosomes are said to be paired when they align along their length and the same genomic locations come in contact with each other. To carry out the pairing, chromosomes have to search for their homologous counterparts in the complex medium inside the nucleus [2, 4, 5]. Once paired, biological events like recombination/crossing over get initiated, and the cells proceed to further stages of meiosis [6]. How cells execute this physical process of recombination/crossing over is still a puzzle.

The fission yeast *Schizosaccharomyces pombe* [7] is a popular model organism to probe chromosome segregation and homologous recombination in different meiotic stages. During early meiosis, an extensive movement of chromosomes has been suggested as the main mechanism of search and pairing of homologous chromosomes [8–15]. This movement is carried out by an extended nuclear oscillation where the whole nucleus is pulled back and forth from one end of the cell to the other. At the beginning of this oscillation, ends of all chromosomes (telomeres) are anchored to a given point known as spindle pole body (SPB) [14, 16–21]. Subsequently, SPB drags the clustered chromosomes for a long duration (*∼* 1 *−* 2 hour), stretching it along the cell axis, before the meiotic division [14, 22–24]. It has been shown that the dragging is carried out by the pulling forces exerted by dynein motors anchored to the cell cortex and microtubules [25–28]. On depleting the dynein motors, the pulling force reduces, thereby impairing the nuclear movement [16, 25, 28]. The movement of SPB along with chromosomes aids the alignment of homologous regions for recombination and also prevents excessive chromosome associations [13]. The elongated nucleus during this movement is called the “horsetail” nucleus and the motion is termed as “horsetail movement” [14, 21, 24, 29–33].

The pioneering experiments to probe the dynamics of homologous pairing during horsetail movement by Ding et al. [34] suggest that the distance between a homologous loci pair decreases during this nuclear oscillation, and the pairing process stops if the oscillation is seized. To examine the dynamics of homologous pairs, different loci were stained and observed under a microscope. It was shown that the association and dissociation of homologous loci is very dynamic, and the frequency of association increases as the horsetail movement goes from stage I to stage V. However, what these stages correspond to and why the frequency of association increases with the progression of stage is not clear.

Even though homologous pairing is a polymer dynamics problem, there have been only a very few theoretical studies that investigate this phenomenon [4, 30, 35–37]. Among these, it is even rarer that someone has probed the horsetail movement dynamics and homologous pairing in *S. pombe* from the polymer dynamics perspective [38, 39]. Lin et al. [38] have performed polymer simulations using a simple model considering the chromosome as a freely jointed chain under external force. Although this model could reproduce results like enhanced contact probability with chromosome movement, it does not explain several features, such as the increasing frequency of association with movement progression. It also lacks information about intra–chromosome interactions that are relevant for chromatin folding/condensation. It is well–known that during cell division chromosomes are in a condensed state [40], probably hindering the alignment and pairing of homologous loci. Hence, it is imperative to investigate the role played by the intra–chromosome attraction on the alignment of homologous pairs. What is the magnitude of force required for this process, and how does it compare with the force needed to open up a folded chromatin are a few interesting questions in the field.

In this work, we investigate the competition between the pulling forces that open up chromatin and the intra–chromatin interaction that folds the chromosome. To achieve this, we employ a bead–spring polymer [41] model for a pair of folded chromosomes connected to SPB in a viscous medium. By creating this *in–silico* version of chromosome and comparing the results of our model to experimental data [34], we not only reaffirm the enhancement of contact probability with loci–dependent pulling, but we also find that during the horsetail movement, different stages correspond to progressive pulling force exerted by the SPB. We show that intra–chromosome attraction impedes the pairing in the early stage of pulling when the polymer is partially open. A subsequent increase in pulling not only leads to amplification in alignment and pairing, it also resolves the non–homologous contact. Further, our results show that the critical force required to unfold the chromatin scales as a power law with the intra–chromatin interaction energy.

## II. MODEL AND METHODS

In the seminal work by Ding et al. [34], recombination among three homologous pairs of chromosomes with different genomic lengths was examined in living cells of *S. pombe* using microscopic techniques. Here, we simulate one such homologous pair of Chr.2 (shown as dark blue and light blue), each having 4.6 Mb length as sketched in Fig. 1(a). Different gene loci studied in the experiment are marked along the chromosome in both pairs by yellow symbols. In the present work, we model the entire loop of each Chr.2 as a coarse–grained linear polymer of *N* = 100 monomers anchored at one end. Modeling looped conformation with linear polymer is reasonable as we are interested in studying the homologous pairing, where what matters the most is the distance from the anchored point [34, 38, 42]. To mimic the clustering of telomere end with SPB, we connect one end of each chain to a larger bead as shown in Fig. 1(b).

**FIG. 1.**
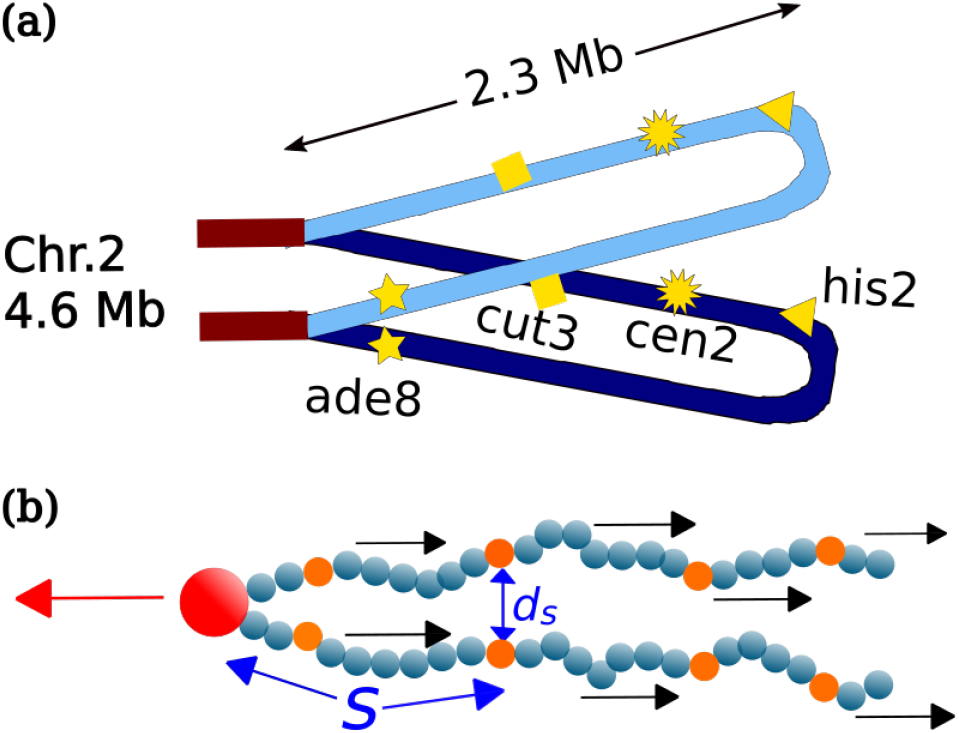
(a) Schematic of a chromosome of genomic length 4.6 Mb used in the experimental work by Ding et al. [34]. Yellow symbols denote various sites on the chromosome that were used to calculate percentage association as a function of distance from the SPB. (b) Schematic of our bead–spring model for a pair of pulled chromosomes. A large red bead represents the SPB, and a pair of chains attached to it represents the homologous chromosome arms. Specific orange beads at different positions of the chain, equidistant from the SPB, are indicative of homologous pairs. *S* represents the contour distance of a bead from the SPB, and *d*_*S*_ is the distance between its homologous pair. The black arrows on the polymer chain denote the relative drag force felt by the monomers when the SPB is pulled in the direction of the red arrow.

In the linear polymer, the neighboring monomers are connected using the harmonic spring potential given by:

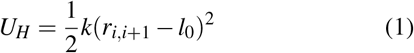

where *r*_*i*,*i*+1_ = |**r**_*i*+1_ *−* **r**_*i*_ |is the distance between consecutive monomers, *k* is the spring constant and *l*_0_ is the equilibrium bond–length. From experiments, we know that chromatin is a folded polymer with loops and intra–chromatin interactions. To account for such attractive interactions, we introduce the Lennard–Jones (LJ) potential:

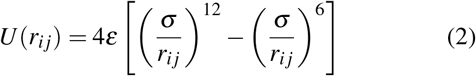

where *ε* is the strength of the interaction and *σ* is the diameter of bead. We have used a cutoff distance *r*_*c*_ = 2.5*σ* and a force correction term that modifies the potential to 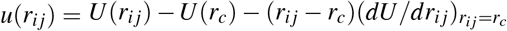[43]. The interaction between the inter-chain monomers as well as the case where there is no attractive interaction within a chain is presented by *u*(*r*_*i j*_ *<* 2^1*/*6^*σ*) = 4*ε*_*w*_[(*σ/r*_*i j*_)^12^ *−* (*σ/r*_*i j*_)^6^ + 1*/*4], the Weeks-Chandler-Andersen (WCA) potential. The interaction of all the monomers with the SPB bead is also considered by the WCA potential.

During the meiotic oscillations, SPB is pulled from one pole of the elongated cell to the other [38]. We mimic this motion by applying a constant pulling force of magnitude *F*_*p*_ = *−βv*_*p*_ on the SPB, where *v*_*p*_ is the speed of the SPB bead. In the comoving frame of the SPB [38, 39], all other monomers experience an equal and opposite force. Hence, the equation of motion for each bead in the under–damped limit is:

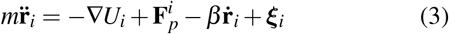

where *m* is the mass of each bead and *U* accounts for the total interaction potential from equations 1 and 2. The drag coefficient *β* is related to the strength of the noise by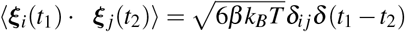, where *k*_*B*_*T* is the thermal energy of the medium. Before applying the pulling force, the system is evolved to the steady state which will form two separate globules around the SPB in case of attractive interaction within a chain. In the next step, we take these states as the initial configurations and apply the pulling force on the monomers.

We fix *k*_*B*_*T* = 0.1, MD time step *δt* = 5 × 10^*−*4^ in all our simulations and vary 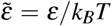 and *F*_*p*_ as the two major parameters. 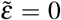 corresponds to the case where there is no intra–chromatin attraction within a chain. We choose *k* = 3000*k*_*B*_*T* to prevent any extension in the equilibrium separation between monomers. The viscous drag coefficient *β* = 1 and *l*_0_ = 1.25*σ*, with *σ* = 1 and time is in units of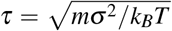. Visualization was done using VMD, and numerical analysis was carried out using custom codes.

### Correspondence with length scales in experiments

To compare our results with the experimental work done by Ding et al.[34], we map the entire loop of Chr.2 (4.6 Mb) to a linear chain with *N* = 100 beads, therefore the contour length of the linear chain is 2.3 Mb (see Fig. 1). As a result, each bead on the polymer corresponds to 23 kb of chromosome, which can be considered to be *≈* 100 *−* 150 nm [44].

## III. RESULTS AND DISCUSSION

### A. Interplay of intra–chromatin interactions and pulling force

Typically, chromosomes in the nucleus tend to be in a collapsed state; therefore, for the homologous pairing to happen, the pulling force that opens up chromatin must overcome the attractive interactions that cause its collapse. To investigate the interplay between these two opposing forces, in Fig. 2(a), we plot the maximum extension of the polymer *r*_*max*_, as a function of the applied force *F*_*p*_, for different values of intra–chromatin attraction strength 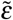. The polymer without any intra–chain attractive interaction, i.e. 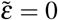, responds to the applied force by a continuous and slow stretching in the direction of the force. However, for any non–zero 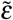, there is a critical pulling force *F*_*c*_, below which there is no significant stretching in the polymer, which is clearly visible as an initial plateau for all 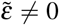. This is an important feature captured in our model, where the strength of intra–chromatin attraction governs the nature of extension due to the applied force. For *F*_*p*_ *> F*_*c*_, the polymer continuously extends and *r*_*max*_ saturates at a value which is independent of 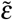, indicating the full extension. The dependence of *F*_*p*_ on 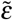 for the full and partial extension of the filament is further elucidated in Fig. 2(b), wherein we plot the 3D distance from SPB (⟨*r*_*S*_⟩) to each genomic location (*S*) along the contour. For very small pulling force (*F*_*p*_ = 0.005), polymer with smaller 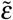 show a partial extension while larger 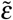 exhibit a plateau at low ⟨*r*_*S*_⟩ for all *S*. The fact that ⟨*r*_*S*_⟩ is independent of *S* for small *F*_*p*_ indicates that the polymer is in a globular state. On increasing *F*_*p*_ we observe an increase in ⟨*r*_*S*_⟩ followed by the flat saturated regions that diminishes, which corresponds to the decrease in the folded globular region. The saturated value of ⟨*r*_*S*_⟩ is same as *r*_*max*_ in Fig. 2(a). Therefore, Fig. 2(a) and Fig. 2(b) show that for an intermediate value of 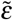 and *F*_*p*_, the polymer exhibits a partially open state with a globule at the end, which stretches completely for large *F*_*p*_. In the open segment of the polymer, ⟨*r*_*S*_⟩ varies linearly with *S*, while in the globule, it remains constant.

**FIG. 2.**
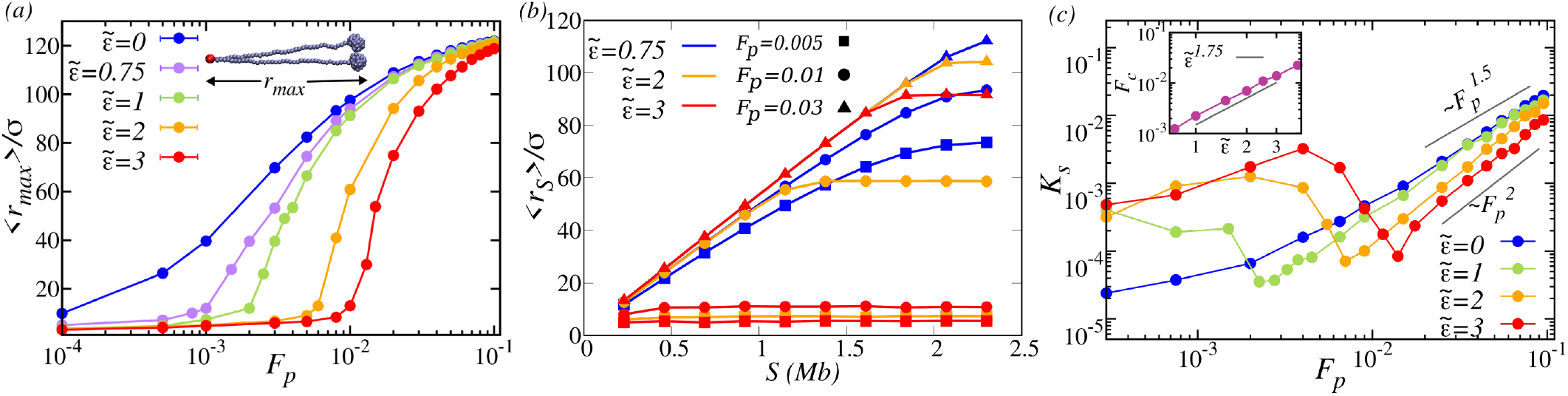
(a) Maximum extension (*r*_*max*_) is plotted against the magnitude of *F*_*p*_ for various values of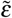. (b) Average distance (⟨*r*_*S*_⟩) of beads from the SPB plotted against genomic location *S*. (c) The stiffness as a function of *F*_*p*_, the exponent for known cases (WLC: 3*/*2, FJC:2) are shown here as a guide to the eye.

The competition between intra–chromatin attractive interaction and the pulling force due to meiotic oscillation is clearly visible in the force–extension curve of Fig. 2(a). This displays a clear transition with a sudden change in *r*_*max*_ as a function of *F*_*p*_. Fig. 2(a) contains a lot of information about the polymer properties. As typically done in pulling experiments, we compute the elasticity (stiffness *K*_*s*_) of the chromatin polymer by calculating the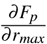 . Fig. 2(c) shows the variation of stiffness (*K*_*s*_) as function of *F*_*p*_. The sudden dip in the stiffness is observed near the critical force *F*_*c*_ that may correspond to the collective opening of the polymer as observed in Fig. 2(a). Beyond this transition, the stiffness shows a power–law dependence as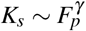. The exponent *γ* in our case lies between the predicted *γ* = 3*/*2 for worm-like-chain (WLC) model and *γ* = 2 for a freely jointed chain (FJC) [45]. Further, the critical force (*F*_*c*_) observed in our simulation may be connected with the experimental observation of the chromatin being in the highly folded and collapsed state before the beginning of the horsetail movement [34]. The inset of Fig. 2(c) shows the variation of *F*_*c*_ with 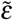that follows the scaling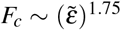. A similar scaling has been reported earlier in the context of collapsed polymer chains undergoing globule–stretch transitions in elongational flow fields [46]. Our studies on the variation of 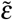 will help experiments to understand how the stretching force will get affected by changing the experimental conditions.

### B. Route to the homologous pairing

During meiosis, the homologous pairs—pairs between the two chains of the chromosome having the same genomic location—come in 3D proximity to copy/exchange the genetic information for the cell division process. The mean separation between the homologous pairs, defined as ⟨*d*_*S*_⟩, is the distance between the beads at the same contour length on the two chains (see Fig. 1). In Fig. 3(a), ⟨*d*_*S*_⟩ is reported for 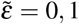 and 3 as a function of *F*_*p*_ for various *S*. As expected, for 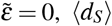 decreases monotonically with increasing *F*_*p*_, indicating the enhancement of homologous pairing with pulling. Further, for a given *F*_*p*_, ⟨*d*_*S*_⟩ increases with *S*, demonstrating that the parts of the chromatin close to the SPB are easily aligned and paired by the pulling force.

**FIG. 3.**
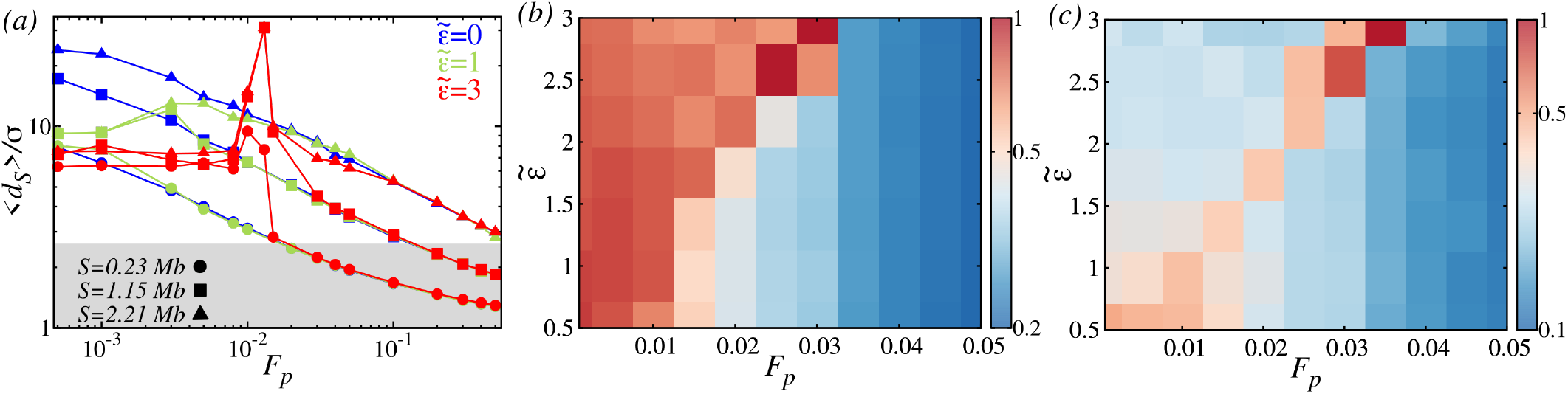
(a) Mean separation (⟨*d*_*S*_⟩) between the homologous pair of beads of the two chains. The shaded region highlights the range of separations between paired homologous beads. (b,c) Color maps of the ⟨*d*_*S*_ ⟩normalized by the maximum value of ⟨*d*_*S*_ ⟩are plotted for *S* = 0.23 and *S* = 1.15 Mb for different values of 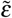 and *F*_*p*_.

Interestingly, for non–zero 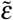, the variation of ⟨*d*_*S*_⟩ is nonmonotonic. At small *F*_*p*_, both the chains remain in a globule state, and hence ⟨*d*_*S*_⟩ is the same for all *S* that corresponds to the mean separation between the center of mass of the two globules. Increasing *F*_*p*_ results in the rise of ⟨*d*_*S*_⟩ until it reaches a maximum; thereafter, it starts to decrease on increasing *F*_*p*_. Note that such non-monotonic behavior is observed for all *S* (see detailed plot in section S1 in the SI [47]), which is further emphasized in the color map in Fig. 3(b) and (c). In the color map, we plot the normalized mean separation ⟨*d*_*S*_⟩ for a particular loci location by varying *F*_*p*_ and 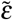. The chosen loci are at *S* = 0.23 Mb (Fig. 3(b)) and *S* = 1.15 Mb (Fig. 3(c)), analogous to the pairs located near and away from the SPB. The non-monotonic behavior with *F*_*p*_ becomes more pronounced for larger 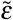 and higher *S*. Therefore, Fig. 3 suggests that the competition between 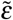 and *F*_*p*_ leads to maximally separated homologous pairs at some intermediate value of *F*_*p*_. A similar behavior is also observed in the variance of the homologous pair separation (See section S1 in the SI [47]).

To comprehend the non–monotonic behavior of ⟨*d*_*S*_⟩ for 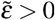, we plot the time evolution of ⟨*d*_*S*_⟩ for different *F*_*p*_ and *S* in Fig. 4(a) for 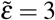. The corresponding representative snapshots are given in Fig. 4(b) (See the corresponding movie in SI[47]). For *F*_*p*_ = 0.008, both the chains remain in the globule state as the pulling force is insufficient, and ⟨*d*_*S*_⟩ is approximately equal to the distance between the center of mass of the globules. For *F*_*p*_ = 0.013, ⟨*d*_*S*_⟩ increases with time and saturates at a large value. This is also consistent with what is observed in Fig. 4(b), where we see the uneven partial opening of either one or both chains. This can be understood if we realize that *F*_*p*_ = 0.013 is close to the critical point of a first order–like sudden transition (See Fig. 2(a)), and one expects huge variability at such transition points [48–50]. On further increasing *F*_*p*_ to 0.05, an initial rise and then a quick decay is observed. The initial growth can again be attributed to the partial opening of the arms and the variability associated with it. Once the chain is completely open, ⟨*d*_*S*_⟩ saturates to smaller values that indicate the homologous pairing. Therefore, for the homologous pairing to happen, starting from a condensed phase of chromatin, the polymer will have to go through the intermediate phase of maximum variability in 3D distance between loci pairs.

**FIG. 4.**
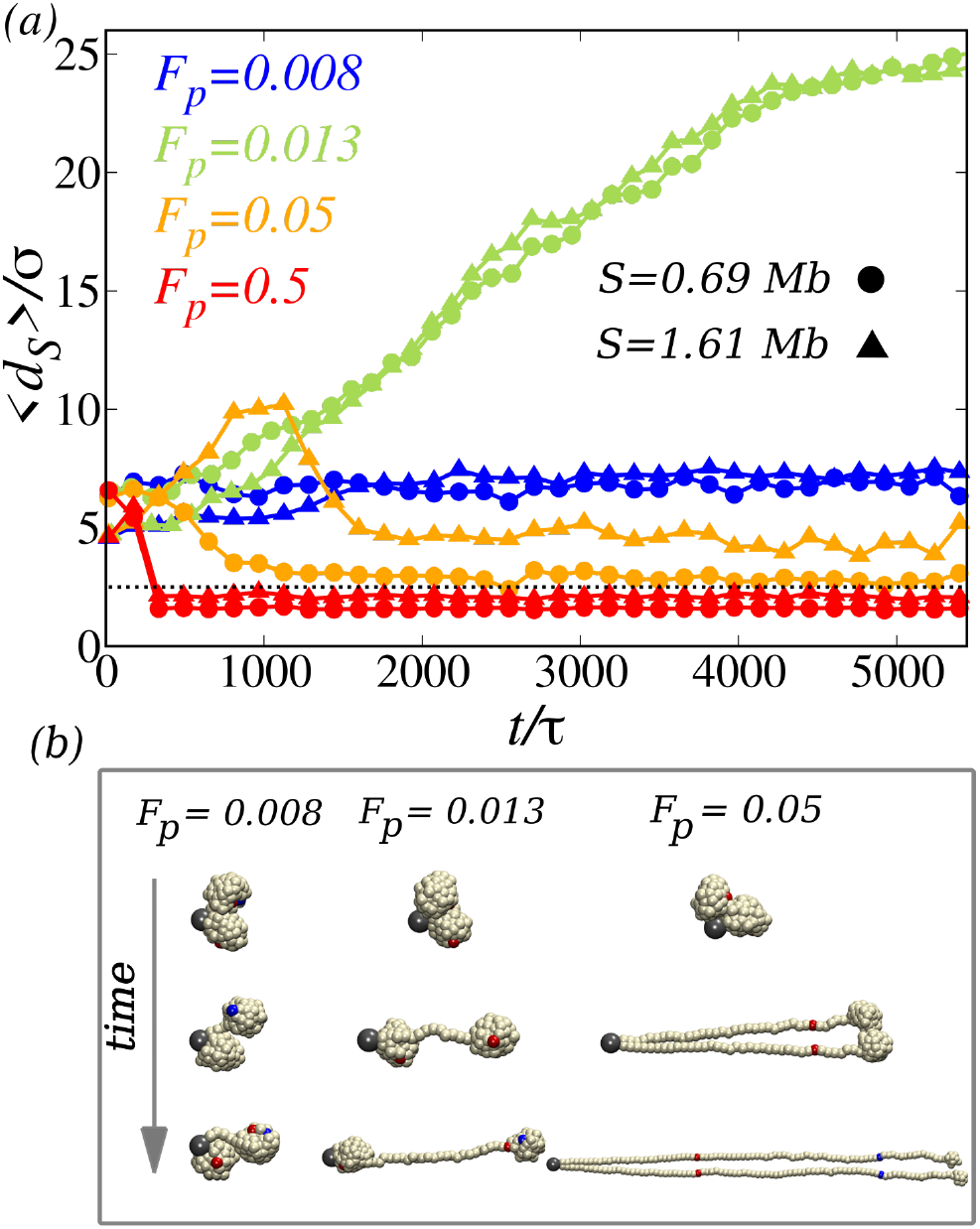
(a) Time evolution of the homologous pair distance for 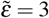 at different *F*_*p*_. Filled circles and triangles correspond to *S* = 0.69 and 1.61 Mb of genomic separation from the SPB. The black dashed line refers to the cutoff distance (*r*_*c*_ = 2.5*σ*) for pairing. (b) Representative snapshots of the modeled chromosome for *F*_*p*_ = 0.008, 0.013 and 0.05.

### C. Mapping of our simulations with experiment: Identification of horsetail stages

The meiotic recombination event in *S. pombe* occurs with the elongation and oscillation of the nucleus across the cell, known as the horsetail movement [34]. In the work of Ding et al. [34], they investigated the spatial configuration of the chromatin during horsetail movement using fluorescent microscopy. They tagged four specific gene loci (on Chr.2) using a GFP marker, which made it possible for them to observe the frequency of association of homologous gene loci when they come in proximity. Our model can be simulated to compute the quantities observed in the experiment precisely. Experiments define two loci to be “associated” when the 3D distance between them is less than 350nm. Consistent with this experimental definition, in our simulations, we consider two homol-ogous loci to be “associated” when the 3D distance between the loci is less than 2.5*σ* .

In the experiments, the entire horsetail movement was divided into five stages, and they observed that the frequency of association increased as the stages progressed. Further, they find that when the genomic distance from SPB to any loci increases, the frequency of association between the corresponding homologous pairs decreases. To understand these observations, we simulate chromatin movement for different applied forces *F*_*p*_, compute contact probability *P*_*c*_ of different homologous loci, and compare them with experiments.

Upon comparison of contact probability (*P*_*c*_) with the genomic distance from SPB (*S*), we find that the different horsetail stages of the experiment can be mapped to different strengths of pulling forces on the SPB. In Fig. 5, we plot *P*_*c*_ from our simulations and the normalized associated loci percentage from experiments. The details of normalization are given in section S2 of the SI text [47]. A remarkable match of horsetail stage I with pulling force *F*_*p*_ = 0.04, stage III with *F*_*p*_ = 0.3 and stage V with *F*_*p*_ = 0.5 is observed. The correspondence between the experiments and our simulation results not only substantiates our model but also suggests that during the nuclear oscillation, different stages of horsetail movements correspond to different magnitudes of pulling forces. Eventhough we have simulated the system with different values of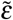, the effect of 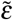 is not prominent in Fig. 5. However, its effect will appear in the subsequent discussions.

**FIG. 5.**
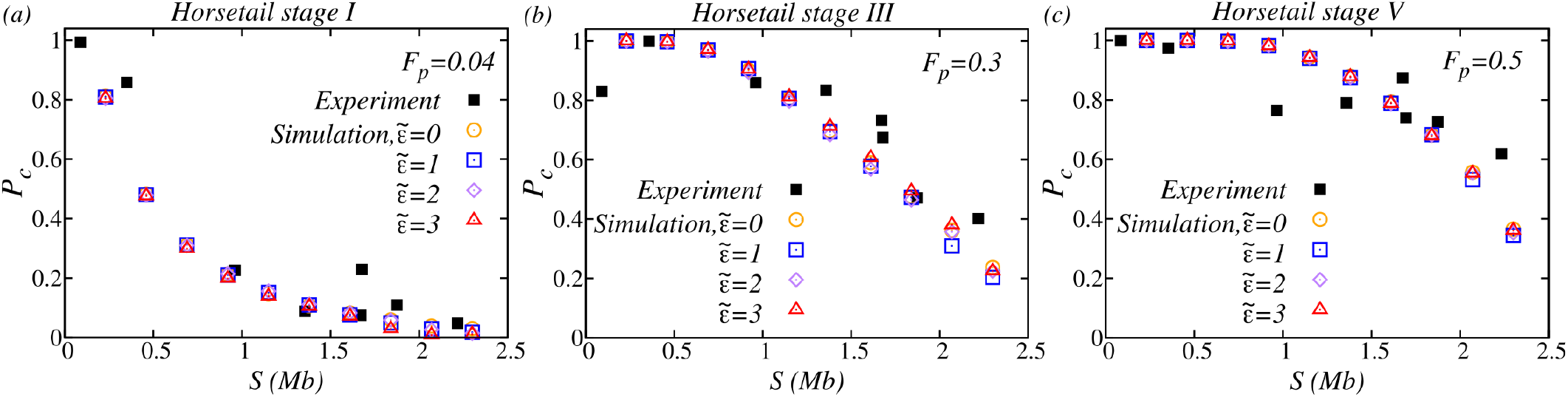
Comparison of the contact probability of homologous loci calculated in our model with the experimental observations [34] at different stages of horsetail movement. (a) Horsetail stage I is compared to *F*_*p*_ = 0.04, (b) Horsetail stage III is compared to *F*_*p*_ = 0.3 and (c) Horsetail stage V is compared to *F*_*p*_ = 0.5.

Identification of horsetail stages I, III and V with *F*_*p*_ = 0.04, 0.3 and 0.5, respectively, allows us to interpolate and associate the remaining stages with appropriate *F*_*p*_. This mapping equips us to further match the frequency of association as a function of the horsetail stages for a given loci pair. Ding et al. have shown that for all loci examined, the loci contact probability increases during the progression of meiotic prophase. For this purpose they have used the loci *ade8* (*S* = 0.35 Mb), *cut3* (*S* = 0.96 Mb), *cen2* (*S* = 1.67 Mb) and *his2* (*S* = 2.21 Mb) of the Chr.2 as shown in Fig. 1(a). To see the effect of SPB proximity on contact probability we plot *P*_*c*_ as a function of *F*_*p*_ for the same loci as given in the experiment. In Fig. 6, a good match between the experiment and our results not only reiterates the increased pairing with the progression of horsetail stages, but also gives more insight into the role of SPB proximity. For the loci closer to SPB (Fig. 6(a) and (b)), *P*_*c*_ increases with *F*_*p*_ and saturates. This shows that the loci sites closer to the SPB are more likely to remain in contact as the horsetail movement progresses to higher stages. Another remarkable feature of Fig. 6 is the scaling of *P*_*c*_ with *F*_*p*_ of different loci. We observe that loci pairs far from the SPB show higher scaling exponent *ν* for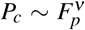. For the *cut3* loci (*S* = 0.96 Mb), *ν* = 1 that changes to *ν* = 1.2 for *cen2* (*S* = 1.67 Mb) and *ν* = 1.5 for *his2* (*S* = 2.21 Mb). This increase in the exponent on increasing the genomic distance of the loci suggests that the loci pairs sitting far away from the SPB have a much stronger dependence on the strength of pulling as compared to loci sitting near SPB. This could be due to the restricted 3D motion of the SPB–proximal loci, which reduces the fluctuations or variance of ⟨*d*_*S*_⟩ . This suggests that although both telomere clustering at SPB and nuclear oscillations are prerequisites for homologous pairing, the latter contributes more to the pairing of homologous loci far from the SPB. Further, note that the effect of 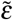 is prominent for smaller values of *F*_*p*_, when these two forces compete against each other. Our results suggest that measuring homologous pairing at higher resolutions during very early stages (earlier than the currently measured stages) will reveal the effect of intra–chromatin interactions 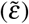 in the experiments.

**FIG. 6.**
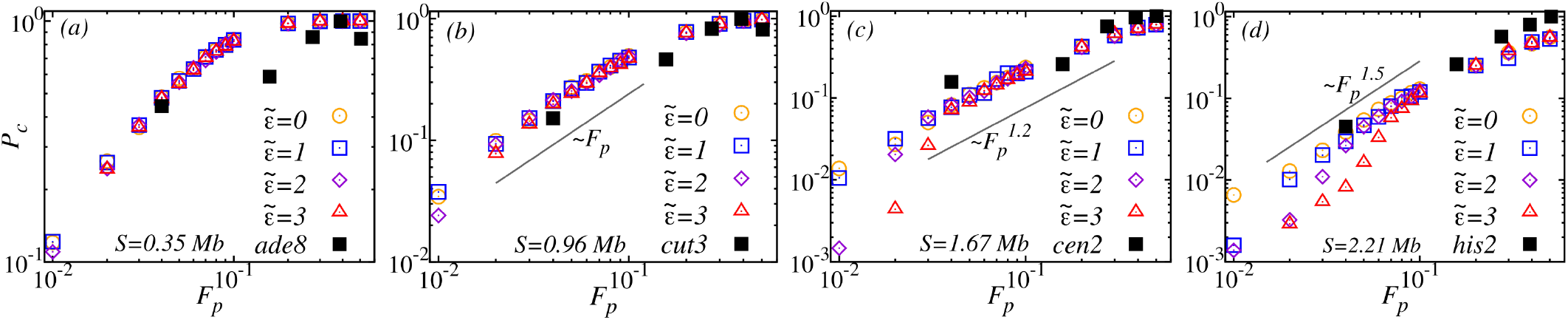
The contact probability between the homologous loci pairs as a function of pulling force *F*_*p*_ for (a) *ade8*, (b) *cut3*, (c) *cen2* and (d) *his2* from the experiments is compared with the loci pair having the corresponding distance in our simulations.

### D. Non–homologous contact

Before the meiotic oscillation, chromatin folding may lead to numerous contacts and interactions between random non-homologous sites. The role of nuclear oscillations during meiosis can also affect the interaction between non–homologous pairs. It was shown by Ding et al. that nuclear oscillations increase the association probability of homologous loci, but no increase was observed for non–homologous loci. In Fig. 7, we plot the probability of contact between a pair of homologous loci (filled symbols) and the probability of contact between a pair of non–homologous loci (unfilled symbols). In principle, the non–homologous loci pair is defined as a pair of beads sitting at different contour distances from the SPB. In this work, we only consider the pairs at bead *i* (at a distance *S*) and bead *i* ± 1 (at a distance *S* ± 0.023 Mb) as the non-homologous pairs. This is because the *P*_*c*_ for a pair whose *S* is very different on two arms is negligible.

**FIG. 7.**
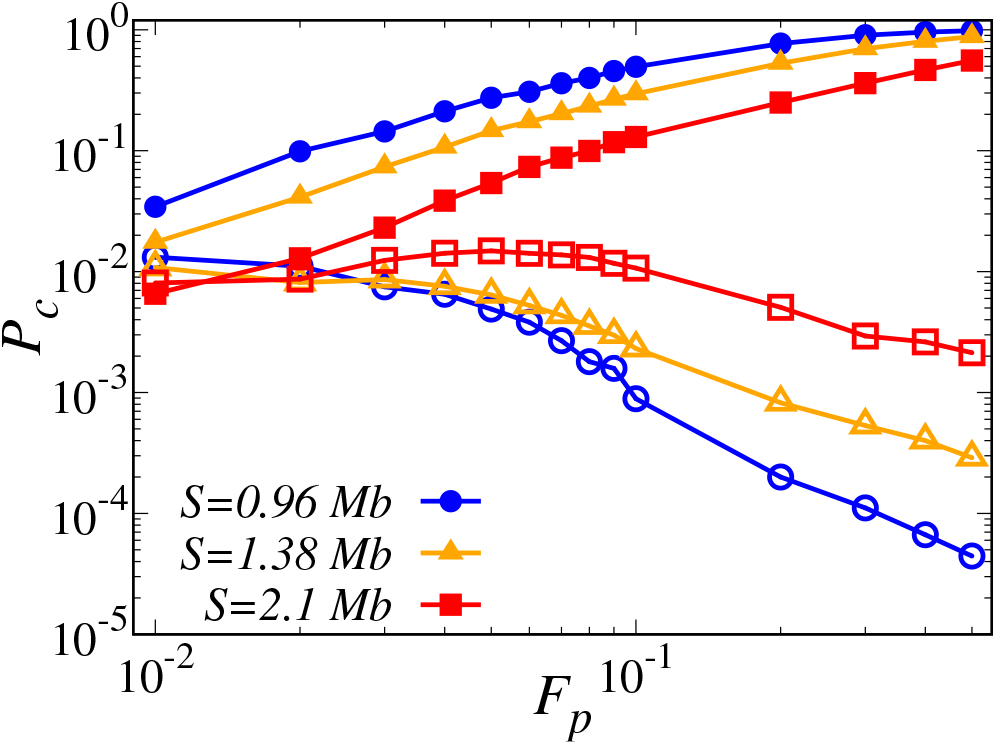
Contact probability of non-homologous adjacent pair compared to homologous pair for 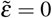 at different contour distance (*S*). Filled and unfilled symbols represent the *P*_*c*_ for homologous and non-homologous pairs, respectively.

The comparison between homologous and non–homologous loci *P*_*c*_ shows that before starting the horsetail stage (*F*_*p*_ = 0.04), the contact probability of both is within the same order of magnitude. However, once the chromosome enters the horsetail stage, the *P*_*c*_ for homologous pairs starts to increase, while for non–homologous pairs, it starts to decrease. This suggests that the force/flow/oscillation has the power to discriminate between different types of pairs, and it can resolve the unwanted interaction and enhance the desired pairing. Such resolution will happen in the initial stage (low *F*_*p*_) of horsetail movement for loci close to SPB (small *S*); however, the loci away from SPB (high *S*) can be resolved only at the later stage (high *F*_*p*_) of the horsetail movement.

## IV CONCLUSION

In this work, we simulate rapid horsetail movement, typically seen during *S. pombe* meiosis, using a polymer model. The existing simulation studies to understand recombination during horsetail movement consider chromatin as an ideal polymer [38]. However, a huge literature of biochemistry experiments has demonstrated that chromatin polymer has an internal structure with several loops mediated by intra–chromatin interactions [51–55]. Further, the single–molecule pulling experiments have also revealed the important role of intra–chromatin interactions [45]. Therefore, here we have investigated how the interplay between the intra–chromatin interactions and pulling forces affect the probability of recombination.

We compare our results with the publicly available experimental data and determine the parameters for which our simulations are realistic. Our results suggest that each stage of the meiotic horsetail movement corresponds to a specific magnitude of pulling force on the SPB identified by a numerical value of pulling force (*F*_*p*_). Next, we attempt to convert them to real units. To do so, we use the equivalence between the non–dimensional representation of *F*_*p*_ in the simulation and real units. Hence, we write

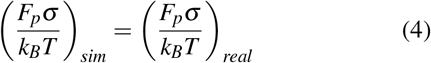

where ()_*sim*_ and ()_*real*_ represent the numerical values of the quantities in simulation and real units, respectively. We take *σ*_*sim*_, (*k*_*B*_*T*)_*sim*_ from the simulation parameters, while *σ*_*real*_ = 100 nm and (*k*_*B*_*T*)_*real*_ = 4.1 pNnm. This suggests that for the horsetail stage I, the *F*_*p*_ = 0.04 corresponds to (*F*_*p*_)_*real*_ = 0.016 pN per monomer. As the SPB drags a pair of chromosomes comprising 200 monomers in our model, the total pulling force applied by the SPB comes out to be *F*_*net*_ = 200 × (*F*_*p*_)_*real*_ = 3.2 pN, in the horsetail stage I. Similarly, for the horsetail stages III and V, the net pulling force exerted by the SPB is estimated to be 24 pN and 40 pN, respectively. Our estimate is in good agreement with the previous work that correlates the pulling forces with the number of dynein motors engaged in pulling the SPB [38].

Due to the limitation of the microscopy techniques, the experimental resolution is typically 350 nm. But the real recombination happens when two segments are in close proximity (a few nm). Our simulations can go beyond the experimentally achieved limit and compute the probability of two regions coming together within the size relevant for realistic recombination. In this work, we show that the effect of intra–chromatin interactions will be more prominent when the pulling forces are low (early stages of pulling) or if the loci is far away from SPB.

## ACKNOWLEDGMENTS

The authors would like to acknowledge the HPC facility at IISER Bhopal for allocating the necessary computational resources. S. T. acknowledges SERB India for research grant number CRG/2022/003778.

