## supplemental_info for "Polymer dynamics reveal stage-wise unfolding and homologous pairing of chromosomes in S. pombe"

#### S1 Results

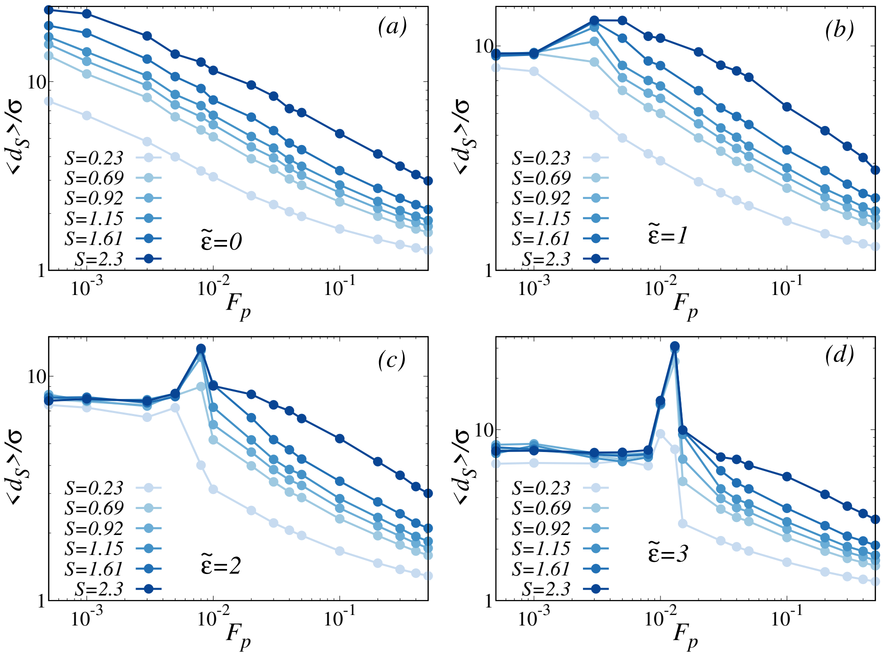


Figure S1: Mean distance ($d_S$) between the homologous pair as a function of pulling force $F_p$ for various $\tilde{\epsilon}$. The figure legend states the value of genomic separation $S$ in Mb.

Detailed plots of the mean separation between homologous gene loci are shown in Fig. S1 for different gene positions. The non–monotonic variation of $\langle d_S \rangle$ becomes more prominent for non-zero $\tilde{\epsilon}$. The same data is plotted in the form of heatmaps for more values of the

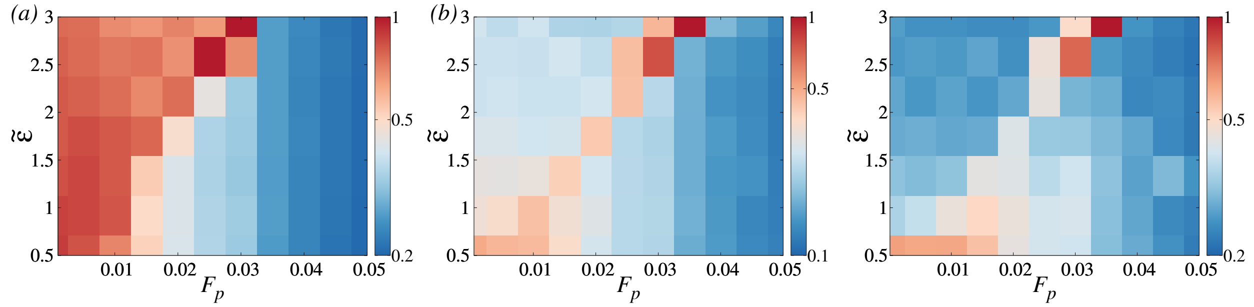


Figure S2: Color map of the mean distance $\langle d_S \rangle$ between homologous pairs calculated at (s) $S = 0.23$ Mb, (b) $S = 1.15$ Mb and (c) $S = 2.3$ Mb at different values of $\tilde{\epsilon}$ and $F_p$.

intra–chromosome interaction strength $\tilde{\epsilon}$ in Fig.S2 and for three different loci pair positions. The standard deviation of $d_S$ is defined as $SD(d_S) = \sqrt{\sum_{i=1}^{n}(\langle d_S \rangle - d_S^i)^2/n}$, where $n$ is the sample size in steady state. In Fig. S3, the standard deviation of $d_S$ is reported separately for different $\tilde{\epsilon}$ at various gene loci $S$. As observed in the mean ($\langle d_S \rangle$) plot in Fig. S1 (and main text), the standard deviation also shows similar non–monotonic behavior for non–zero $\tilde{\epsilon}$. Similar to the color map plotted for the mean ($\langle d_S \rangle$), we also plot the color map of the normalized standard deviation for three different $S$ in Fig. S4. The non–monotonic behavior is clearly observed for higher $\tilde{\epsilon}$ and larger distances from the SPB.

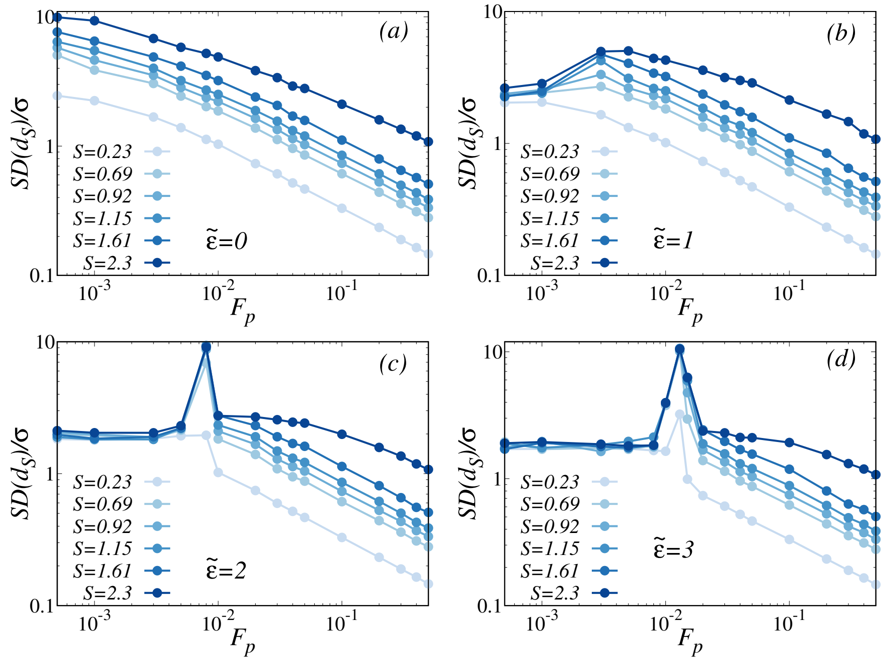


Figure S3: Standard deviation ($SD$) of $d_S$ from mean value for various homologous pairs as a function of pulling force $F_p$ for various $\tilde{\epsilon}$. The figure legend states the value of genomic separation $S$ in Mb.

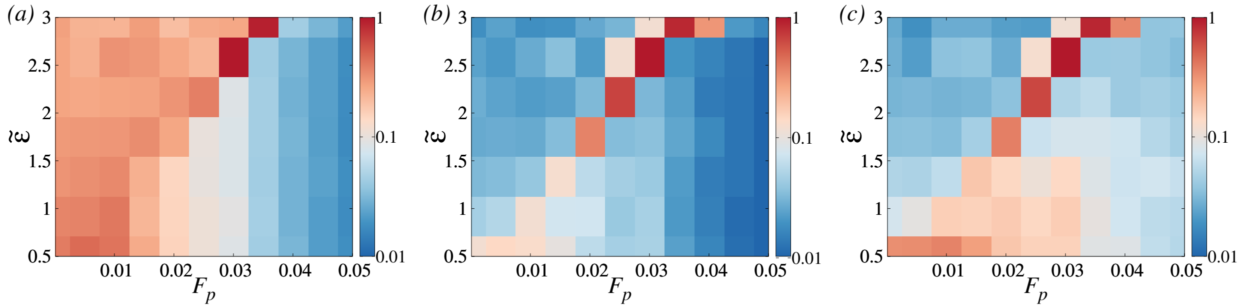


Figure S4: Color map of the standard deviation of $d_S$ calculated for (a) $S = 0.23$ Mb, (b) $S = 1.15$ Mb and (c) $S = 2.3$ Mb at different values of $\tilde{\epsilon}$ and $F_p$.

### S2 Normalization details

The associated loci percentage defined in the experiments by Ding et al. [1] is similar to the contact probability between homologous loci defined in our simulations. To have a direct comparison of the experimental data with our results and for better representation, we normalize the associated loci percentage (or contact probability) in the following way. In Fig. 5 of the main manuscript, all the $P_c$ values of the experimental data were normalized by their highest value taken separately in the three plots. By doing so, the y-axis of the experimental data starts from 1 just like the simulation data. A similar normalization approach was chosen in Fig. 6.

### S3 Movie description

Movie.mp4 video shows an example of the trajectory followed by the modeled chromosome pair for pulling force $F_p = 0.008, 0.013$ and $0.05$; corresponding to the plot and snapshots in Figure. 4 in the main text. Two different homologous loci pair at $S = 0.69$ and $1.61$ Mb are highlighted in black and blue color beads respectively, and the SPB is shown as a large red bead.
